# Serial enrichment-based quantitative proteomics enables deep and multiplexed profiling of post-translational modifications

**DOI:** 10.64898/2026.01.15.697874

**Authors:** Mohd. Altaf Najar, Anjana Aravind, T. S. Keshava Prasad, Prashant Kumar Modi

## Abstract

Post-translational modifications (PTMs) play central roles in regulating protein function, localization, stability, and signaling networks. Comprehensive characterization of multiple PTMs, however, remains technically challenging due to limited sample availability, enrichment, incompatibilities, and analytical complexity. Here, we present an integrated serial enrichment workflow that enables quantitative profiling of multiple PTMs from a single biological sample. Using a streamlined strategy that combines StageTip-based fractionation with sequential immunoaffinity and metal oxide affinity enrichment, we achieved robust identification and quantification of lysine acetylation, lysine succinylation, phosphotyrosine, and global phosphorylation within the same sample. Application of this workflow resulted in deep proteome coverage alongside high-confidence PTM site identification with minimal sample loss and high reproducibility. Importantly, serial enrichment preserved PTM specificity and enabled comparative quantitative analysis across modification types. This approach provides a practical and scalable solution for integrated multi-PTM analysis, facilitating comprehensive interrogation of proteome regulation under limited sample conditions and offering broad applicability to systems biology, disease profiling, and translational proteomics studies.

## Introduction

Post-translational modifications (PTMs) of proteins constitute a fundamental regulatory layer that expands proteome complexity and enables precise control over protein structure, localization, stability, and activity [1–3]. More than 300 distinct PTMs have been described to date, including phosphorylation, ubiquitination, acetylation, methylation, glycosylation, and SUMOylation, among others [4]. These modifications dynamically regulate nearly all cellular processes, such as signal transduction, transcriptional regulation, cell-cycle progression, DNA damage response, metabolism, and immune signaling [5, 6]. Dysregulation of PTM landscapes is strongly associated with numerous human diseases, including cancer, neurodegenerative disorders, metabolic syndromes, and inflammatory conditions, highlighting the importance of comprehensive PTM characterization [1, 7].

Mass spectrometry (MS)-based proteomics has emerged as an unparalleled analytical platform for the identification and quantification of PTMs at a proteome-wide scale [8, 9]. Advances in high-resolution MS instrumentation, peptide separation technologies, and computational analysis pipelines have enabled the detection of thousands of modified peptides in a single experiment [10]. Despite these technological developments, PTM analysis remains challenging due to the typically low stoichiometry of modified peptides, their transient and dynamic nature, and the overwhelming abundance of unmodified peptides, which suppress PTM signals during MS acquisition. Consequently, selective enrichment strategies are essential to enhance PTM detection sensitivity and depth of coverage.

Most PTM-focused proteomic studies traditionally concentrate on a single modification type, employing dedicated enrichment methods such as immobilized metal affinity chromatography (IMAC) or metal oxide affinity chromatography (MOAC) for phosphopeptides, antibody-based enrichment for ubiquitinated or acetylated peptides, and lectin-based approaches for glycopeptides [11, 12]. While these targeted strategies have yielded invaluable insights into specific PTM-driven regulatory mechanisms, they fail to capture the intricate crosstalk and coexistence of multiple PTMs on the same proteins or within the same biological system. Since PTMs frequently act in a coordinated or antagonistic manner, a holistic, multi-PTM analytical approach is required to fully understand cellular regulation at the systems level.

Recent efforts have attempted to address this limitation by developing workflows for parallel or sequential enrichment of multiple PTMs from a single biological sample. However, many existing approaches suffer from practical limitations, including high sample input requirements, workflow complexity, incompatibility between enrichment chemistries, and loss of material during multiple handling steps. These challenges restrict their applicability, particularly when working with limited or precious biological samples such as clinical specimens.

In this study, we present a streamlined and efficient strategy for the serial enrichment of multiple PTMs from a single peptide sample. Our workflow integrates sequential enrichment steps optimized to preserve sample integrity while enabling selective isolation of distinct PTM classes. By coupling serial PTM enrichment with robust peptide fractionation using StageTip-based separation, this approach maximizes proteome coverage and PTM depth without increasing sample input requirements. This methodology enables comprehensive interrogation of PTM landscapes and provides a scalable platform for studying PTM interplay in complex biological systems.

## Results

To evaluate the impact of serial PTM enrichment and StageTip-based fractionation on proteome and PTM-ome depth, we applied a sequential enrichment workflow to a TMT-labeled peptidome (figure 1). The serial enrichment strategy targeted lysine succinylation, lysine acetylation, phosphotyrosine, and global phosphorylation using Fe-NTA and TiO₂ enrichment approaches. To systematically assess the contribution of enrichment and fractionation to PTM identification, multiple sample types were analyzed. These included an un-enriched (UE) proteome sample, generated from 5% of the input material prior to enrichment and fractionated using C18 StageTips; an unfractionated (UF) sample, consisting of 10% of each enriched fraction analyzed without further fractionation; and a fractionated (FR) sample, in which the remaining 90% of each enriched fraction was subjected to C18 StageTip-based fractionation. In addition, the final flow-through (FT) remaining after all enrichment steps was collected, of which 10% was fractionated using C18 StageTips and analyzed.

**Figure 1.**
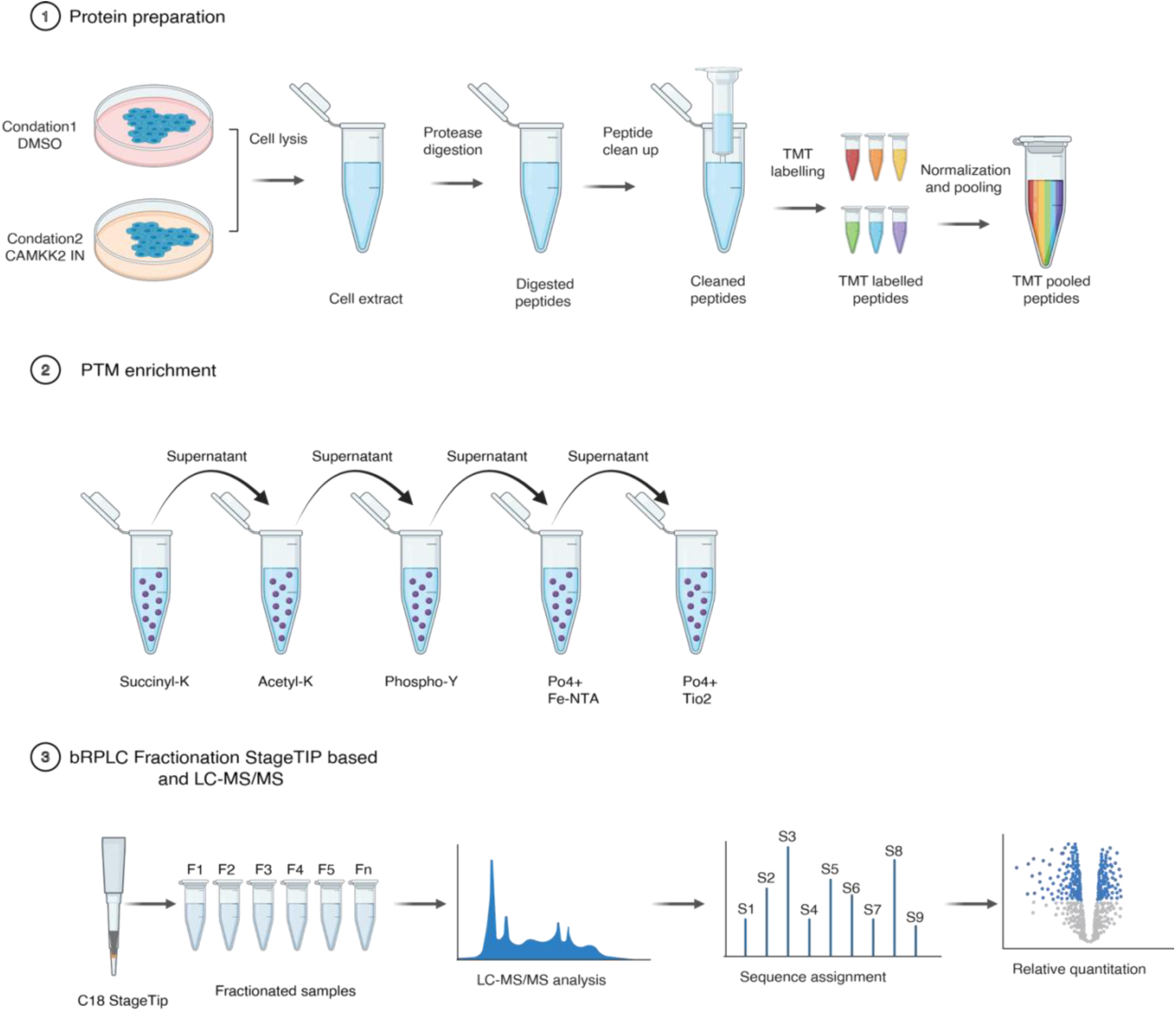
Overview of the serial PTM enrichment–based quantitative proteomics workflow. **(1)** Cells were lysed, reduced, alkylated, and digested to generate tryptic peptides, which were quantified, TMT-labeled, and pooled. **(2)** The combined peptide mixture was subjected to a serial enrichment strategy targeting lysine succinylation, lysine acetylation, phosphotyrosine, and global phosphorylation, with flow-through fractions retained at each step. **(3)** Enriched PTM fractions and selected flow-through and total proteome samples were further fractionated using C18 StageTips prior to LC–MS/MS analysis, enabling deep and multiplexed quantitative profiling of multiple PTMs from a single biological sample.

Using this workflow, we quantified 5,600 proteins and identified approximately 10,696 phosphopeptides across phosphotyrosine immunoaffinity purification, Fe-NTA, and TiO₂ enrichment strategies. Among these, 6,659 phosphopeptides were identified following Fe-NTA enrichment, and 9,114 phosphopeptides were identified by TiO₂ enrichment. Phosphotyrosine-specific analysis revealed 683 peptides harboring tyrosine phosphorylation. In addition to phosphorylation, serial PTM enrichment enabled the identification of 10,064 succinylation sites through succinylome analysis and 2,996 acetylation sites from acetylome enrichment.

Serial PTM enrichment, combined with StageTip-based fractionation, substantially enhanced coverage of both the proteome and the PTM-ome. StageTip fractionation effectively partitioned peptide complexity across multiple fractions, thereby reducing sample complexity and minimizing ion suppression during mass spectrometric analysis. As shown in Figure 2, fractionation resulted in a marked increase in the number of identified peptides and PTM sites compared with unenriched or unfractionated samples, underscoring the benefit of sequential enrichment coupled with offline peptide separation for comprehensive PTM profiling.

**Figure 2.**
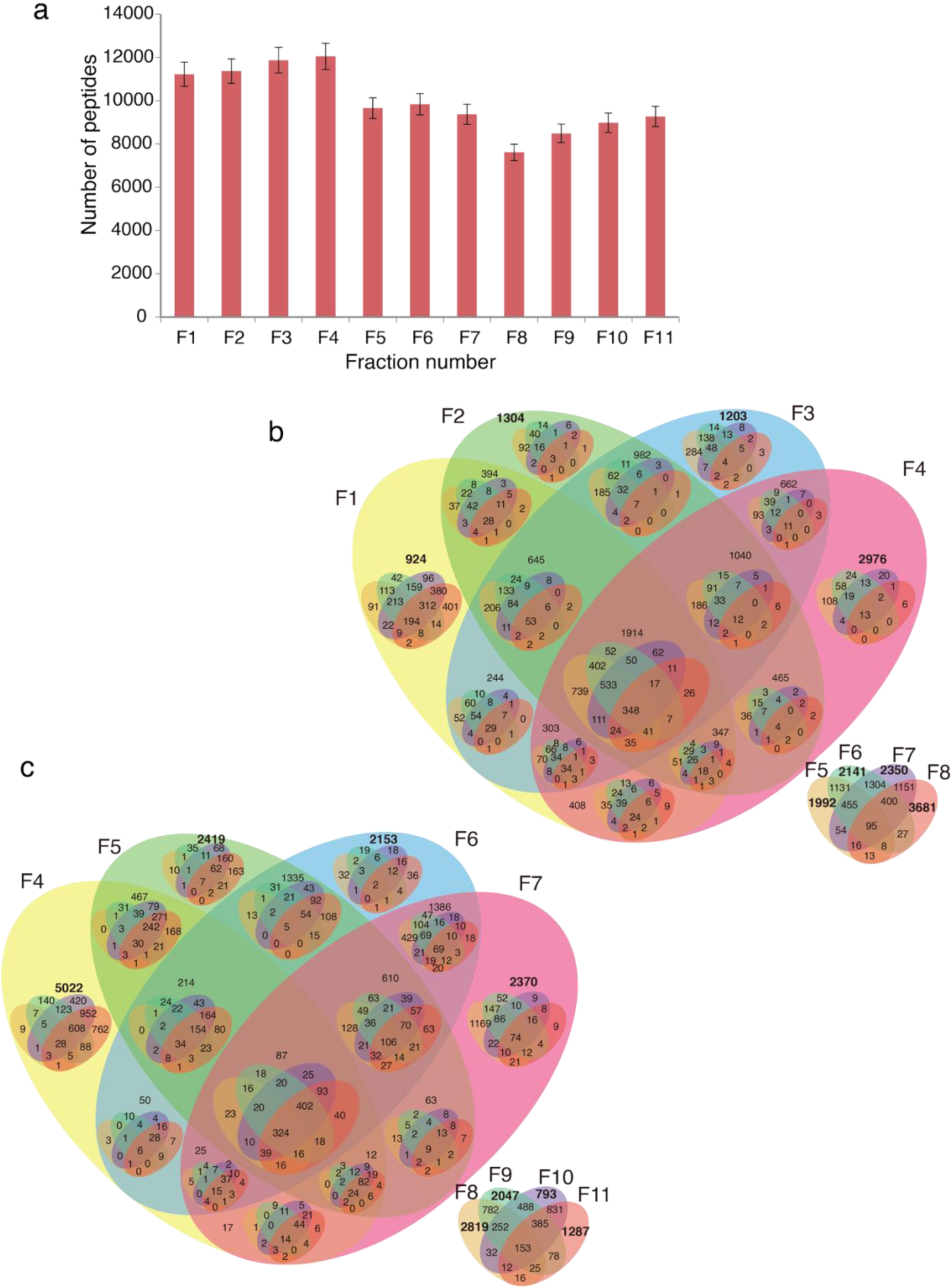
StageTip-based fractionation enhances proteome and PTM-ome depth through complementary peptide identification across fractions. **(a)** Bar plot showing the distribution of identified peptides across individual StageTip fractions (F1–F11). Fractionation resulted in relatively balanced peptide recovery across fractions, indicating effective partitioning of peptide complexity prior to LC–MS/MS analysis. **(b, c)** Venn diagram analyses illustrating peptide overlap and uniqueness across StageTip fractions for enriched samples. A substantial proportion of peptides were uniquely identified in individual fractions, with limited redundancy between neighboring fractions, demonstrating orthogonal peptide separation and efficient reduction of sample complexity.

Importantly, analysis of peptide overlaps across individual fractions revealed that a substantial proportion of peptides and PTM-modified sites were uniquely identified in specific fractions, with limited redundancy between neighboring fractions. Venn diagram analyses demonstrate that each fraction contributed a distinct subset of peptides, indicating efficient orthogonal separation and minimal carryover between fractions. This fraction-specific peptide distribution highlights the ability of the StageTip-based strategy to expand analytical depth by capturing low-abundance and otherwise masked PTM species that would be missed in single-shot analyses. Collectively, these results demonstrate that offline fractionation not only increases total identification numbers but also enhances the diversity and coverage of the PTM landscape by enabling complementary sampling across fractions.

To quantitatively assess enrichment efficiency, we compared the number of modified peptides identified across unenriched (UE), unfractionated enriched (UF), flow-through (FT), and fractionated enriched (FR) samples. As illustrated in Figure 3, enrichment alone increased PTM identification relative to UE samples; however, the combination of enrichment and fractionation (FR) yielded the highest number of modified peptides across all PTM classes analyzed. This trend was consistently observed for acetylation, phosphorylation (Fe-NTA and TiO₂), phosphotyrosine, and succinylation, underscoring the broad applicability and robustness of the serial enrichment–fractionation workflow.

**Figure 3.**
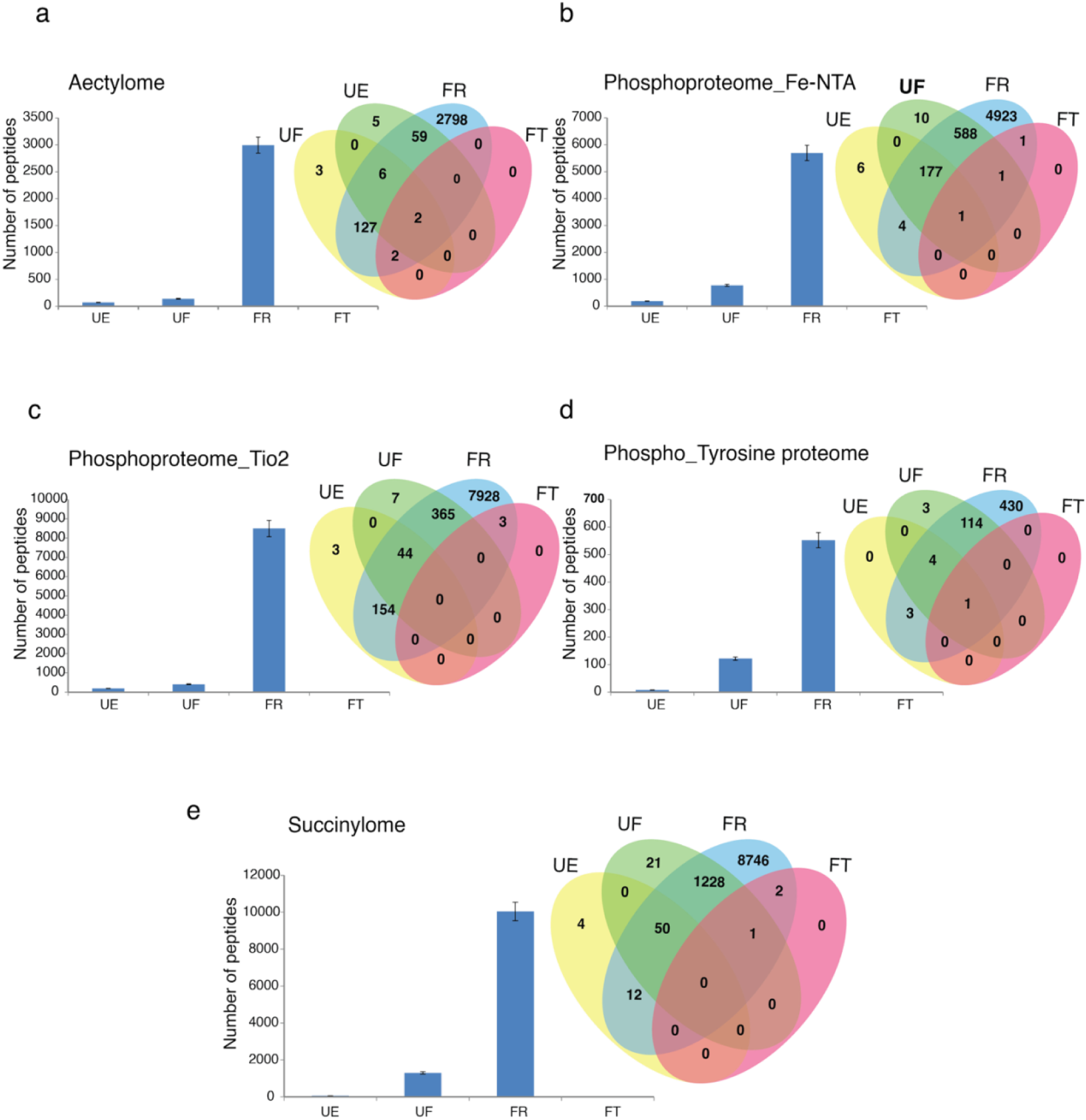
Impact of serial PTM enrichment and StageTip fractionation on PTM identification depth. Bar plots depict the total number of identified modified peptides, and Venn diagrams illustrate the overlap and uniqueness of PTM sites across different sample processing strategies for **(a)** acetylome, **(b)** phosphoproteome enriched by Fe-NTA, **(c)** phosphoproteome enriched by TiO₂, **(d)** phosphotyrosine proteome, and **(e)** succinylome analyses. Samples include unenriched peptides (UE), enriched but unfractionated peptides (UF), enriched and C18 StageTip-fractionated peptides (FR), and the final flow-through (FT) collected after serial enrichment. Across all PTM classes, enrichment substantially increased PTM detection compared with unenriched samples, whereas StageTip fractionation further enhanced coverage by enabling the identification of numerous unique PTM-modified peptides distributed across fractions. These data demonstrate that serial enrichment combined with offline peptide fractionation markedly improves PTM-ome depth and complexity resolution.

Serial enrichment enabled the identification of both PTM-unique and PTM-shared peptides (Figure 4a). For phosphorylation, three complementary enrichment strategies were employed, allowing near-comprehensive recovery of phosphopeptides from the proteome. In total, 10,696 phosphorylation sites were identified, including an overlap of 59 phosphotyrosine sites detected across all three enrichment approaches. The phosphotyrosine immunoaffinity purification assay yielded approximately 443 unique phosphotyrosine-containing peptides. Fe-NTA enrichment isolated 5,711 phosphopeptides, including 1,735 unique sites, 3,900 sites shared with TiO₂ enrichment, and 17 phosphotyrosine sites overlapping with phosphotyrosine immunoenrichment. TiO₂-based enrichment identified the largest number of phosphopeptides, comprising 4,506 unique sites in addition to 3,900 shared with Fe-NTA and 36 shared with phosphotyrosine enrichment (Figure 4b).

**Figure 4.**
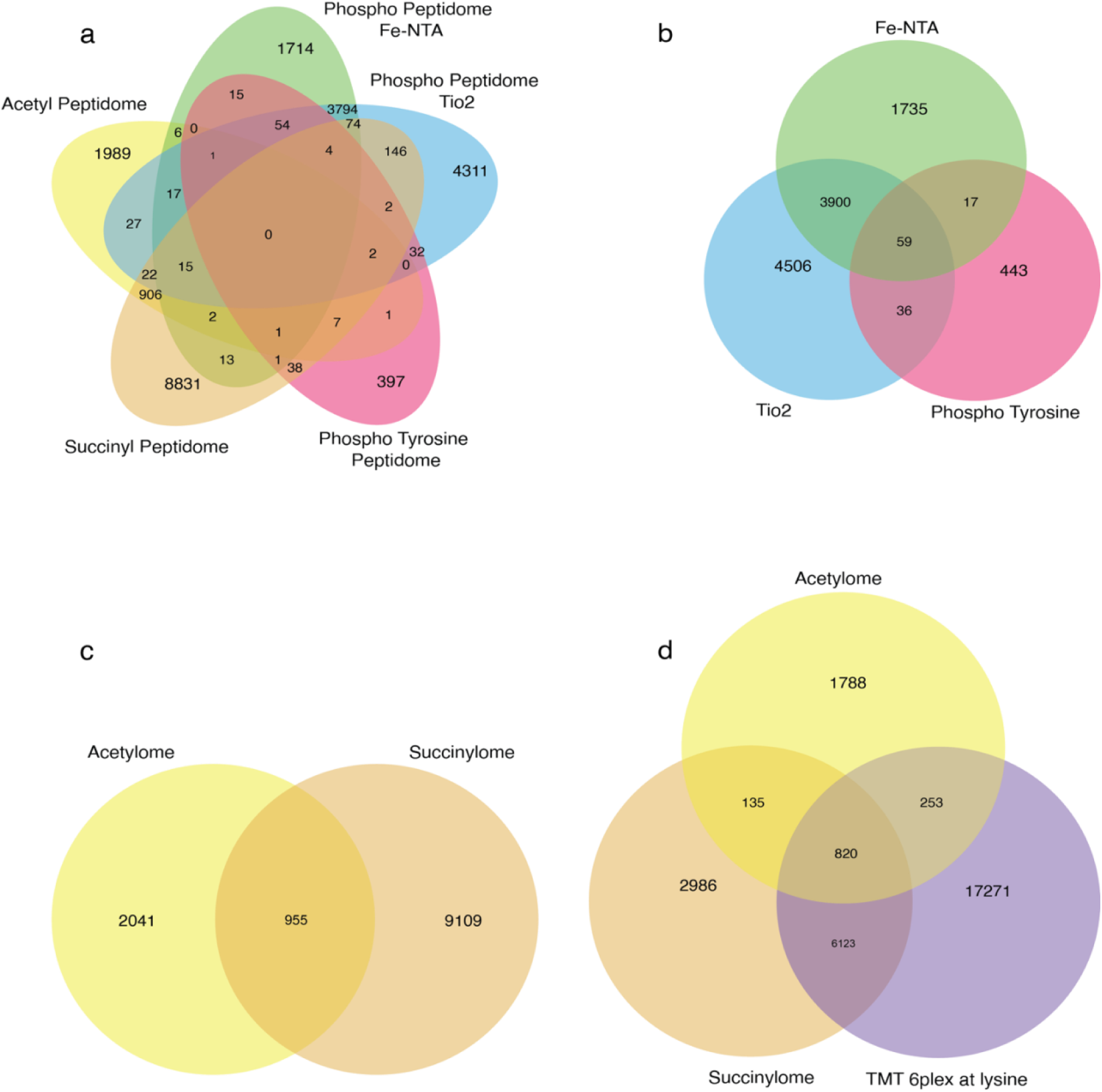
Overlap and complementarity of PTM identification enabled by serial enrichment. Venn diagrams illustrate the overlap and uniqueness of PTM-modified peptides identified using the serial enrichment workflow. **(a)** Overlap of phosphopeptides identified across three complementary phosphorylation enrichment strategies, including phosphotyrosine immunoaffinity purification, Fe-NTA enrichment, and TiO₂ enrichment, demonstrating the complementarity of these approaches for comprehensive phosphoproteome coverage. **(b)** Pairwise and shared overlap of phosphotyrosine, Fe-NTA–enriched, and TiO₂-enriched phosphopeptides, highlighting both unique and commonly identified phosphorylation sites across enrichment methods. **(c)** Overlap between lysine acetylated and lysine succinylated peptides, revealing extensive crosstalk between lysine acylation PTMs. **(d)** Overlap among acetylated, succinylated, and TMT-modified peptides, reflecting shared lysine targeting while confirming the identification of both PTM-specific and uniquely modified peptide populations within the serial enrichment workflow.

In addition to phosphorylation, crosstalk between lysine acetylation and succinylation PTMs was observed. A total of 955 peptides were identified harboring both acetylation and succinylation modifications (Figure 4c). Because acetylation, succinylation, and TMT labeling all target lysine residues, both shared and unique peptides carrying these modifications were detected (Figure 4d). To assess whether TMT labeling interferes with succinylation site identification, we examined peptides shared between TMT-modified and succinylated datasets. Manual inspection of MS/MS spectra revealed distinct immonium ions corresponding to succinylated and TMT-modified lysine residues (Figures 5b and 5d). Furthermore, characteristic MS² fragmentation signatures unique to succinylated and TMT-modified lysines were clearly distinguishable (Figures 5a and 5c), confirming that TMT labeling does not compromise the confident identification of succinylation sites.

**Figure 5.**
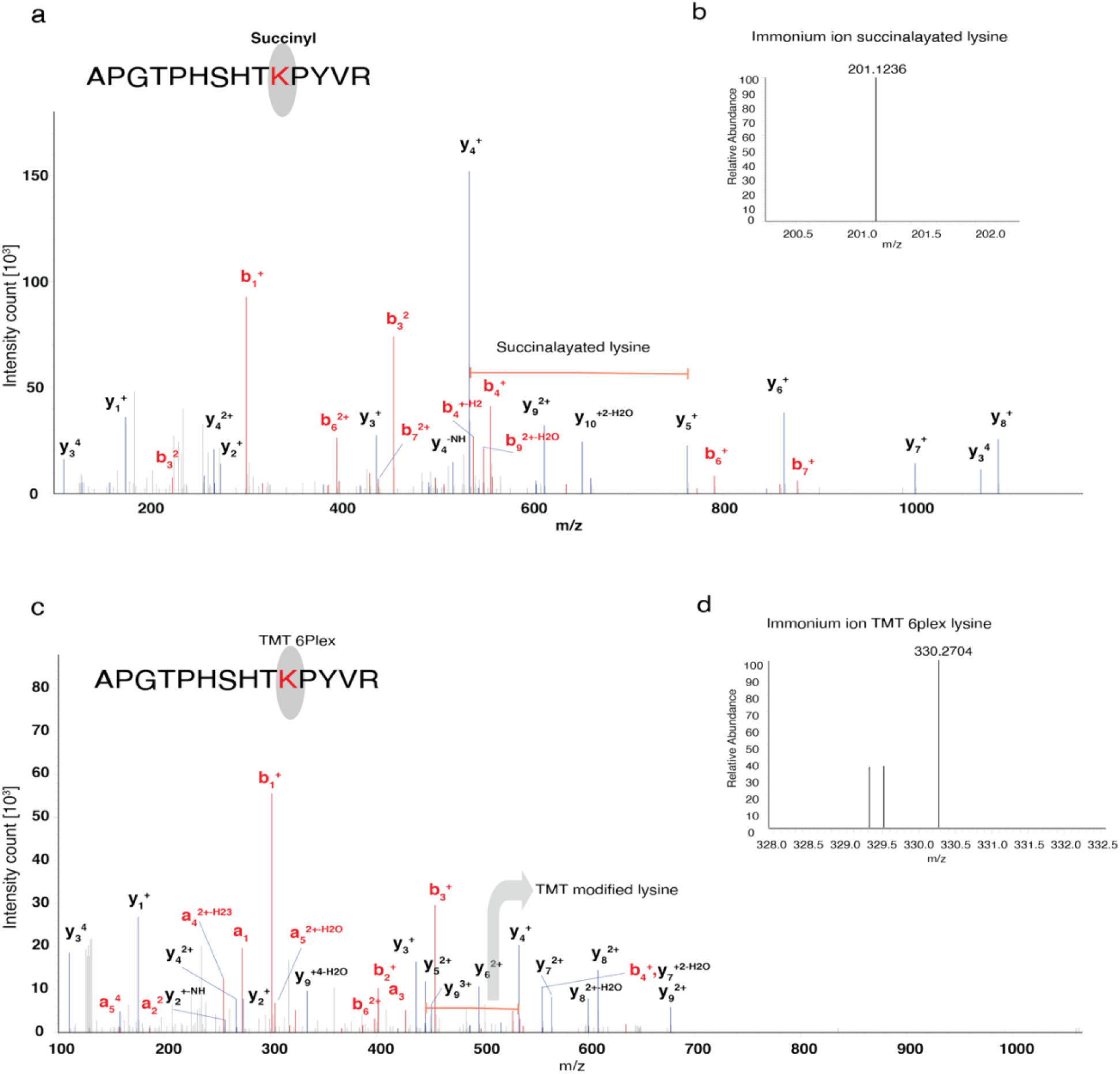
Distinct MS/MS fragmentation signatures enable confident discrimination of succinylated and TMT-modified lysine residues. Representative MS/MS spectra of peptides harboring lysine succinylation and TMT modification are shown. **(a)** MS² spectrum of a succinylated peptide highlighting characteristic fragment ions and neutral loss patterns specific to lysine succinylation. **(b)** Expanded low-mass region of the MS² spectrum showing diagnostic immonium ions corresponding to succinylated lysine. **(c)** MS² spectrum of a TMT-modified peptide displaying fragmentation features characteristic of TMT-labeled lysine residues. **(d)** Expanded low-mass region of the MS² spectrum highlighting TMT-specific reporter and immonium ions. Distinct fragmentation patterns and diagnostic ions observed for succinylated and TMT-modified peptides demonstrate that TMT labeling does not interfere with the confident identification of lysine succinylation sites within the serial PTM enrichment workflow.

To further evaluate the biological fidelity of the serial PTM enrichment workflow, we examined phosphorylation changes in samples treated with a CAMKK2 inhibitor compared to vehicle control. Consistent with our previous findings demonstrating CAMKK2-dependent activation of the MEK/ERK signaling axis, phosphoproteomic analysis revealed widespread hypophosphorylation of proteins involved in cell-cycle regulation and DNA replication upon CAMKK2 inhibition [12] (Figure 6). Notably, reduced phosphorylation was observed on key replication licensing and mitotic regulators, including minichromosome maintenance complex components (MCM2 and MCM3), the cell-cycle regulator CDC20, and components of the MAPK signaling pathway. These phosphorylation changes are consistent with the established role of CAMKK2-mediated Ca²⁺/calmodulin signaling in promoting proliferative signaling via MEK/ERK and downstream cyclin-dependent kinase activity. Importantly, the concordance between the phosphoproteomic data generated using the serial enrichment strategy and previously reported CAMKK2-regulated signaling pathways underscores the robustness and biological relevance of the workflow for capturing physiologically meaningful PTM dynamics.

**Figure 6.**
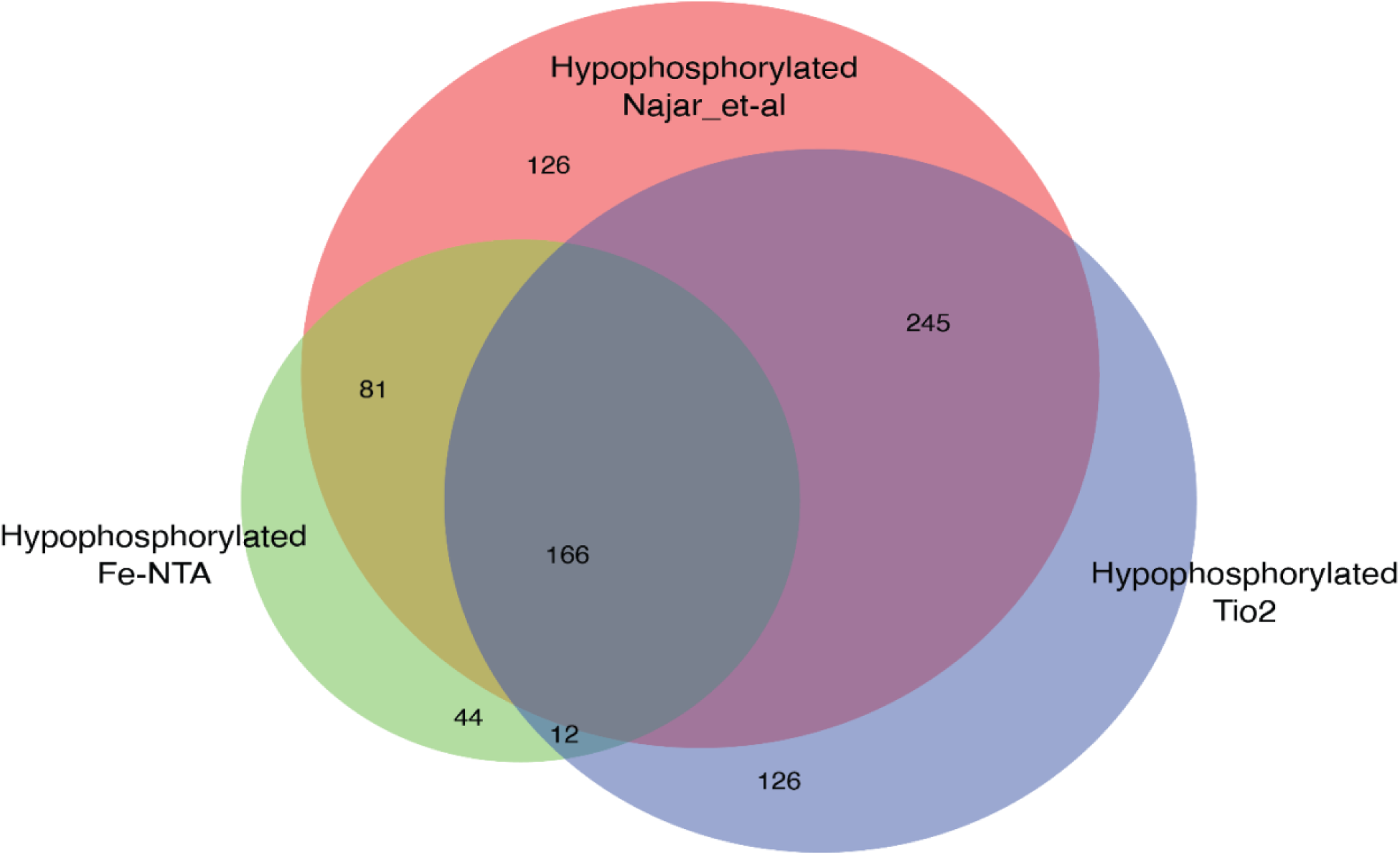
Concordance of CAMKK2 inhibition–induced hypophosphorylation identified by serial PTM enrichment with previously reported CAMKK2-regulated signaling. Venn diagram showing the overlap of hypophosphorylated proteins identified in the current phosphoproteomic analysis following CAMKK2 inhibitor treatment with hypophosphorylated proteins previously reported in CAMKK2-dependent signaling Najar *et al.* [12] Hypophosphorylated proteins identified using Fe-NTA and TiO₂-based phosphopeptide enrichment strategies are shown, highlighting shared and method-specific phosphorylation changes. The substantial overlap across datasets includes key regulators of DNA replication, cell-cycle progression, and MAPK signaling, supporting the biological fidelity and robustness of the serial PTM enrichment workflow in capturing physiologically relevant phosphorylation dynamics.

## Methods

### Preparation of samples of AGS cells for quantitative PTM-ome

AGS cells were obtained from the American Type Culture Collection (ATCC, Manassas, VA) and maintained in Dulbecco’s Modified Eagle Medium (DMEM, high glucose; Gibco) supplemented with 10% dialyzed fetal bovine serum (FBS; Gibco), 5% sodium bicarbonate, and 1× penicillin/streptomycin (Gibco). Cells were cultured in a humidified incubator at 37 °C with 5% CO₂. For quantitative PTM-ome analysis, cells were seeded and allowed to adhere and grow for 24 h under both control and treatment conditions. Subsequently, cells were exposed to the CAMKK2 inhibitor for a total duration of 48 h. Immediately prior to harvesting, cells were treated with kinase and deacetylase inhibitors for 15 min to preserve endogenous phosphorylation and acetylation states. All experiments were performed in biological triplicates. Following protein extraction and digestion, peptides from each condition were labeled using TMT 6-plex reagents according to the manufacturer’s instructions. Labeled samples corresponding to biological triplicates were then combined in equal amounts and pooled for downstream fractionation and mass spectrometric analysis.

### Cell lysis and peptide digestion

Following drug treatment, cells were washed three times with ice-cold phosphate-buffered saline (PBS) to remove residual media and inhibitors and harvested by gentle scraping in ice-cold lysis buffer. Cell pellets were lysed for 30 min on ice in urea lysis buffer containing 8 M urea, 75 mM NaCl, 50 mM Tris-HCl (pH 8.0), 1 mM EDTA, and phosphatase inhibitor cocktail (Thermo Scientific, 78441), with intermittent vortexing to ensure efficient solubilization. Lysates were clarified by centrifugation at 13,000 × g for 30 min at 4 °C, and the resulting supernatants were collected for downstream processing. Protein concentration was determined using the bicinchoninic acid (BCA) assay (Thermo Fisher Scientific, 23225).

For proteolytic digestion, protein disulfide bonds were reduced with 5 mM dithiothreitol (DTT; Sigma-Aldrich, D9779-25G) for 30 min at 60 °C, followed by alkylation with 20 mM iodoacetamide (IAA; Sigma, I6125-10G) for 30 min in the dark at room temperature. To facilitate enzymatic digestion, samples were diluted with 50 mM triethylammonium bicarbonate (TEABC; pH 8.0) to reduce the urea concentration to 1 M. Proteins were then digested overnight at 37 °C with sequencing-grade trypsin (Promega, V5111) at an enzyme-to-substrate ratio of 1:20 under constant agitation. Digestion was quenched by acidifying the samples with 0.1% formic acid (Fluka Analytical, 94318-250ML-F). Digestion efficiency was assessed by resolving 10 µg of the peptide digest on SDS-PAGE.

Peptide mixtures were desalted using C18 Sep-Pak cartridges (Waters, 500 mg, WAT036790). Columns were conditioned with 5 mL of 100% acetonitrile and equilibrated with 3 × 5 mL of 0.1% trifluoroacetic acid (TFA). After sample loading, columns were washed with 3 × 5 mL of 0.1% TFA followed by a final wash with 5 mL of 1% formic acid. Peptides were eluted with 3 mL of 50% acetonitrile containing 0.1% formic acid. Eluted samples were dried in a vacuum concentrator to remove organic solvent and obtain purified peptide mixtures.

For downstream mass spectrometry analysis, 10% of the total peptide material was reserved for unfractionated global proteome analysis, while the remaining peptides were subjected to PTM-specific enrichment workflows.

### TMT Labeling

Prior to TMT labeling, peptide concentrations were quantified using the Pierce™ Quantitative Colorimetric Peptide Assay (Thermo Fisher Scientific, 23275) according to the manufacturer’s instructions to ensure accurate normalization across samples. TMT labeling was performed based on a previously described protocol [13], with minor modifications.

For labeling, equal amounts of peptides (500 µg per sample) were used. Peptide samples were reconstituted in 400 µL of 50 mM triethylammonium bicarbonate (TEABC, pH 8.0). TMT 10-plex Mass Tag labeling reagents (Thermo Scientific, 90110) were equilibrated to room temperature for 5 min prior to use and dissolved in 40 µL of anhydrous acetonitrile (Sigma-Aldrich) immediately before labeling.

For this study, six TMT channels were employed. TMT labels 126, 127, and 128 were assigned to vehicle-treated control samples, whereas labels 129, 130, and 131 were used to label CAMKK2 inhibited samples. The labeling reactions were carried out for 1 h at room temperature with gentle mixing. Following incubation, reactions were quenched by the addition of 8 µL of 5% hydroxylamine and incubated for an additional 15 min at room temperature. To assess labeling efficiency and verify equal peptide representation across channels, 10 µL aliquots from each labeled sample were collected prior to pooling and analyzed separately.

After confirming labeling efficiency, equal amounts (450 µg) of TMT-labeled peptides from each channel were combined into a single pooled sample. The pooled sample was subsequently dried in a vacuum concentrator and stored at −80 °C until further fractionation and PTM enrichment.

### K(Succinyl), K(Acetyl) and Y(Phospho) enrichment of peptides by immuno-affinity purification

Succinyl, acetyl, and phosphotyrosine peptide enrichment was done using the PTMScan kit for respective PTMs (Cell Signaling Technology, 13416, 13764, 38572). Enrichment was performed according to the manufacturer’s protocol, as described in the previous paper [11]. Each enrichment was done in a serial fashion flowthrough of first enrichment was used for second enrichment as shown in second section of figure 1. Before enrichment, the peptide sample was divided into two equal aliquots. Each aliquot was adjusted to a final volume of 1.4 mL with immuno-affinity purification (IAP) buffer and mixed thoroughly to fully dissolve the peptides. After dissolving the peptides pH was checked and peptide mixture was centrifuged at 10,000 g for 10 minutes at 4^0^C, the clear supernatant was taken in a separate microcentrifuge tube and cooled in ice. For washing the antibody beads, the beads were first centrifuged at 2000 g for 30 seconds at 4^0^C to remove the storage buffer and then washed with 1 mL of 1X PBS 4 times, and finally the beads were resuspended in 40 µL of 1X PBS. The clear peptide mixture was transferred to a microfuge tube containing antibody beads and incubated on a rotator at 4^0^C for 2 hours. After incubation the beads were pelleted down at 2000 g for 30 seconds and supernatant was collected for the next enrichment. The beads enriched with peptides were washed with 1 mL of IAP buffer twice followed by wash with 1 mL ice cold LC-MS grade water (Merck) 4 times. Finally, the enriched peptides were eluted with 0.15 % of TFA, for that 55 µL of 0.15% TFA were added on the beads and incubated at room temperature for 10 minutes and then centrifuged at 2000 g, supernatant was collected and beads were resuspended in elution solution and repeated the step. The supernatant collected after the enrichment step was used for the next PTM enrichment and it was carried out in a serial way.

### IMAC phospho-enrichment of peptides using Fe-NTA beads

Enrichment of phosphopeptides was done using the Fe-NTA phosphopeptide enrichment kit (Thermo Fisher Scientific, A32992). Enrichment was done according to the manufacturer’s protocol. The flow-through collected after immunoaffinity purification was dried in a vacuum concentrator and resuspended in the binding buffer provided in the Fe-NTA kit. Two phases were observed: an immiscible layer at the bottom and a clear, miscible layer at the top. The top portion was taken for the next step. Washing of Fe-NTA beads was started by the removal of storage buffer, followed by washing with 50 µL washing buffer three times at 2000 g. A clear peptide mixture was transferred to the column and incubated for 30 minutes at room temperature, and after every 5 minutes, the beads were mixed with gentle tapping of the column. Washing of unbound peptides was done with 100 µL wash buffer at 2000 g for 30 seconds twice, and a final wash with LC-MS grade water. Finally, peptides were eluted with 50 µL elution buffer at 2000g twice. Eluted phosphopeptides were then dried using a vacuum concentrator.

### IMAC phospho-enrichment of peptides using TiO_2_ beads

Flow-through from phospho Fe-NTA enrichment was first dried in a vacuum concentrator, then resuspended in DHB solution (5% 2,5-dihydroxybenzoic acid, 80% ACN, 3% TFA, HPLC grade) and allowed to dissolve completely in a vortex mixer. TiO_2_ beads (GL Sciences, 5020-75010) were taken in a ratio of 1:2 (bead: peptide). The required amount of beads was taken in a microfuge tube and beads were dehydrated in a dry bath at 95^0^C for 10 minutes keeping the lid open and then beads were let to cool at room temperature. Finally, beads were resuspended in 400 µL of DHB solution and incubated at room temperature for 15 minutes on the rotator. The peptide mixture was then split equally in 4 microfuge tubes. Equal volume of beads was added in each sample vial. Samples were then incubated at room temperature on the rotator for 1 hour after incubation beads were spin down at 2000 g for 1 minute and supernatant was collected and dried in vacuum concentrator. Further, washing was done using the wash solution 1 (80% ACN, 3% TFA, HPLC grade). 1 mL of wash solution was added to the beads, and the solution was gently mixed by inverting the tube 10 times. The wash solution was removed by centrifugation at 2000 g for 1 minute. Next wash was given by wash solution 2 (80% ACN, 1% TFA, LC-MS grade water). For the third wash, 100 µL of wash solution 3 (80% ACN, 0.1% TFA, LC-MS grade water) was added, and the entire content was transferred to the StageTip with a C8 plug. The wash solution was removed by centrifugation at 2000 g for 5 minutes; this step was repeated by adding 100 µL of wash solution 3. Finally, phospho-peptides were eluted in a microfuge tube containing 10 µL of 3% formic acid, which was kept on ice. For elution, 80 µL of elution solution (4% NH4OH, LC-MS grade water) was added and eluted at 2000 g at 4^0^C for 5 minutes, the elution was repeated twice. Eluted phospho-peptides were then dried in a vacuum concentrator and stored in −80^0^C until further use.

### Basic reversed-phase liquid chromatography (bRPLC) separation of peptides using C18 StageTip

To reduce peptide complexity, samples were separated by basic reversed-phase chromatography using C18 filled StageTip column. Unlike other pre-fractionation techniques, basic reversed-phase separation enables the simultaneous generation of equal and distinct fractions of modified and unmodified peptide mixtures. Post-translational modifications such as phosphorylation, succinylation and acetylation bias the length and charge state of peptides. It is found that separation techniques that rely on the charge state or pKa value of peptides, such as SCX or IEF, do not provide the level of uniqueness/fraction nor the equivalency of modified peptide numbers/fraction obtained using basic reversed-phase separation with non-contiguous fraction combining. Further fractionation may be of value in improving PTM identification, at the cost of increased instrument time. The extent of peptide fractionation was tailored to sample complexity, with the highest level of fractionation applied to highly complex samples such as the total proteome and enrichment flow-through prior to phosphoproteome analysis, and reduced fractionation used for less complex PTM-enriched samples, including Fe-NTA– and TiO₂-based phosphoproteomics, as well as succinylation, acetylation, and phosphotyrosine proteome analyses. Further, using C18 StageTip, we have done fractionation of enriched samples. The usage of StageTip will reduce the sample loss and can be used in any lab without having an offline HPLC facility.

For column preparation, C18 material (Empore, 66883-U) was used; 10 mg of the C18 material was taken and packed in a T200 tip (Axygen T-200-C). Column charging was done by adding 100 µL of ACN (LCMS grade) and removed the solvent at 2000 g for 5 minutes, followed by column equilibration by passing 100 µL of 0.2% trifluoroacetic acid at 2000 g for 5 minutes, both column charging and equilibration was repeated twice. Enriched, proteome and flow-through peptides were reconstituted in 100 µL of 0.2% TFA, pH 2.0 and centrifuged at 10,000 g to clarify the mixture before it was loaded on to the column. Sample loading was carried out at 1500 g for 15 minutes at 25^0^C and was repeated thrice. After sample loading, washing was done with 100 µL 0.2% TFA and was repeated thrice followed by peptide separation.

Peptide separation was done using mobile phase-A (10 mM TEABC in LC-MS grade water) and mobile phase-B (100% ACN LC-MS grade) by centrifugation at 2000 g and 25^0^C. Mobile phase A and B was mixed in a separate micro centrifuge tube for each fraction as shown in table 1and then loaded on the column and eluted at 2000 g in separate tubes labelled as fraction (1,2,3,4……24). The samples were fractionated in 24 fractions and then serpentine, concatenated pattern was followed for combining eluted fractions from the beginning, middle and end of the run to generate sub-fractions of similar complexities that contain hydrophilic as well as hydrophobic peptides. For proteome, flowthrough, phosphoproteome Fe-NTA enriched, succinylome were combined into 12 fraction the fractions were combined (1,13; 2,14; 3,15; 4,16……12,24), and acetylome, phospho tyrosine, phospho TiO_2_ DHB enriched samples were combined into, 4,8 and 6 fractions respectively. The fractions were vacuum dried and stored at −20^0^C before MS analysis.

**Table 1.** Mobile phase composition used for StageTip C18–based peptide fractionation. Stepwise elution conditions for StageTip C18 fractionation are shown, indicating the acetonitrile concentration (%) and corresponding volumes of mobile phase A and mobile phase B used for each fraction. Peptides were sequentially eluted using increasing acetonitrile concentrations to achieve effective fractionation prior to LC–MS/MS analysis.

| **Fraction number** | **Acetonitrile (%)** | **Volume mobile phase A (μL)** | **Volume mobile phase B (μL)** |
| --- | --- | --- | --- |
| 1 | 4 | 96 | 4 |
| 2 | 5 | 95 | 5 |
| 3 | 6 | 94 | 6 |
| 4 | 7 | 93 | 7 |
| 5 | 8 | 92 | 8 |
| 6 | 9 | 91 | 9 |
| 7 | 10 | 90 | 10 |
| 8 | 11 | 89 | 11 |
| 9 | 13 | 87 | 13 |
| 10 | 15 | 85 | 15 |
| 11 | 17 | 83 | 17 |
| 12 | 19 | 81 | 19 |
| 13 | 21 | 79 | 21 |
| 14 | 23 | 77 | 23 |
| 15 | 25 | 75 | 25 |
| 16 | 27 | 73 | 27 |
| 17 | 29 | 71 | 29 |
| 18 | 31 | 69 | 31 |
| 19 | 33 | 67 | 33 |
| 20 | 35 | 65 | 35 |
| 21 | 37 | 63 | 37 |
| 22 | 40 | 60 | 40 |
| 23 | 45 | 55 | 45 |
| 24 | 50 | 50 | 50 |

### LC-MS/MS analysis

All peptide samples were separated on an online nanoflow EASY-nLC 1200 UHPLC system (Thermo Fisher Scientific) and analyzed on an Orbitrap Fusion Tribid mass spectrometer (Thermo Fisher Scientific). Peptide digests were reconstituted in 0.1% formic acid and loaded onto the trap column (75 μm × 2 cm) at a flow rate of 3 μl/min. Injected peptides were separated at a flow rate of 300 nL/min with a stepped gradient 120 min gradient from 2% solvent A (0.1% formic acid) to 30% solvent B (80% acetonitrile, 0.1% formic acid), 10 min 70% solvent A (0.1% formic acid) to 60 % solvent B (80% acetonitrile, 0.1% formic acid), 6 min 60 % solvent A (0.1% formic acid) 100% solvent B (80% acetonitrile, 0.1% formic acid) 4 min 100 % 100% solvent B (80% acetonitrile, 0.1% formic acid). Data-dependent acquisition was obtained using Xcalibur software using NSI ion source in positive ion mode at a spray voltage of 26.00 kV, ion transfer tube temperature 275 ^0^C. MS1 Spectra were measured with a resolution of 1,20,000 an AGC target of 2e^5^ and a mass range from 400 to 1600 m/z with a maximum injection time of 50 ms. MS2 data was acquired in top speed with a duty cycle of 3 sec at 60,000 resolution, an AGC target of 1.0e^5^, HCD mode was used for fragmentation with a stepped collision energy of 35 and ± 3 %. Scan range mode was set as Auto: m/z normal with starting m/z from 100, and maximum injection time of 200 ms.

### Identification and quantification of proteins

All mass spectra were analyzed with Proteome Discoverer software version 2.2 using Sequest HT and Mascot search engine against human RefSeq database version 109. MS/MS searches for the proteome data sets were performed with the following parameters: Oxidation of methionine and protein N-terminal acetylation as variable modifications; carbamidomethylation as fixed modification; TMT 6plex as fixed modification; phosphorylation of serine, threonine and tyrosine residues; succinylation at lysine and epsilon-acetylated lysine was used for total proteome and flow-through samples. Phosphorylation of serine, threonine and tyrosine residues along with oxidation of methionine and protein N-terminal acetylation as variable modifications; carbamidomethylation and TMT 6plex as fixed modification for immune affinity of phospho-Y, IMAC Fe-NTA and TiO_2_ enriched samples. Epsilon-acetylated lysine, oxidation of methionine and protein N-terminal acetylation as variable modifications; carbamidomethylation as fixed modification; TMT 6plex as fixed modification for K(Ac) enriched samples. Succinylation for K(Su) and oxidation of methionine and protein N-terminal acetylation as variable modifications; carbamidomethylation as fixed modification; TMT 6plex as fixed modification enriched samples. Trypsin was selected as the digestion enzyme, and a minimum of 7 amino acids and 2 missed cleavages per peptide were allowed. Minimum precursor mass was set as 350 Da and maximum precursor mass as 8000 Da. The mass tolerance for precursor ions was set to 20 ppm and fragment mass tolerance was set to 0.02 Da. For identification, we applied a maximum FDR of 1% separately on protein, peptide, and PTM-site levels. The ptmRS node was used for PTM localization. phosphoRS was set as false so that all the PTMs other than phospho will be located along with phospho modification. Reporter ion quantifier node was used to quantify the TMT reporter ion across the samples. PTM-sites were fully localized when they were measured with a localization probability >0.75 in each of the three replicates.

## Discussion

Comprehensive profiling of post-translational modifications (PTMs) remains analytically challenging due to their low stoichiometry, chemical diversity, and the limited compatibility of enrichment strategies within a single workflow [14]. Most PTM-focused proteomic studies rely on parallel enrichment pipelines, requiring large sample inputs and increasing experimental variability [14, 15]. To our knowledge, phosphopeptide enrichment has not been previously integrated as a serial step following immuno-affinity–based PTM enrichment within a unified workflow [16]. In this study, we establish a serial PTM enrichment strategy combined with TMT-based quantification and StageTip fractionation that enables deep and multiplexed profiling of multiple PTMs from a single biological sample using both IAP and metal affinity enrichments.

A major advantage of this workflow is its efficient use of sample material. Sequential enrichment of succinylation, acetylation, and phosphorylation from a single TMT-labeled peptidome enabled identification of more than 10,000 phosphorylation and succinylation sites, nearly 3,000 acetylation sites, and over 5,600 proteins. These results demonstrate that serial enrichment does not substantially compromise downstream PTM recovery and provides coverage comparable to PTM-specific workflows, while significantly reducing sample consumption and experimental redundancy.

StageTip-based fractionation proved to be a critical determinant of proteome and PTM-ome depth. Fractionated samples consistently yielded substantially higher numbers of modified peptides compared to unfractionated or unenriched controls. By reducing peptide complexity prior to LC–MS/MS analysis, fractionation improved sampling efficiency and enabled detection of low-abundance PTM species. Importantly, enrichment alone was insufficient to achieve maximal coverage, highlighting the complementary role of fractionation even in PTM-enriched samples.

Integration of complementary phosphopeptide enrichment strategies further enhanced phosphoproteome coverage. Fe-NTA, TiO₂, and phosphotyrosine immunoaffinity enrichment displayed overlapping yet distinct selectivity profiles, resulting in recovery of both shared and unique phosphorylation events. The combination of these approaches enabled broad capture of serine/threonine phosphorylation while selectively enriching low-abundance phosphotyrosine peptides, demonstrating the value of multi-chemistry enrichment within a unified workflow.

Lysine acylation analysis revealed extensive overlap between acetylation and succinylation sites, underscoring the importance of workflows capable of capturing multiple acyl modifications simultaneously. Because TMT labeling also targets lysine residues, potential interference between isobaric labeling and lysine acylation was a key concern. Manual inspection of MS/MS spectra confirmed distinct fragmentation signatures and immonium ions for TMT-modified and succinylated lysines, validating the compatibility of TMT-based quantification with serial lysine acylation enrichment.

While serial enrichment increases experimental complexity, careful optimization of enrichment order, sample handling, and fractionation mitigated potential losses and variability. The workflow is readily scalable and compatible with multiplexed quantitative proteomics, making it suitable for studies in which sample availability is limited or integrated PTM profiling is required.

In conclusion, this study establishes a robust and scalable serial PTM enrichment strategy that enables deep, quantitative profiling of multiple PTMs from a single sample. By integrating TMT labeling, complementary enrichment chemistries, and StageTip fractionation, this workflow provides an efficient platform for multi-PTM proteomics and offers a practical alternative to parallel, PTM-specific analytical pipelines.

## Ethics approval and consent to participate

Not applicable

## Data Availability Statement

All the data related to the study are provided in the manuscript and supplementary data.

## Author Contributions

P. K. M. conceived the idea, designed the experiments and interpretation of data, and critically reviewed and edited the manuscript. M.A.N conceived the idea, designed the experiments along with P.K.M. performed experiments and data analysis, drafted the manuscript, and prepared figures. A.A. performed data analysis and drafted the manuscript. T.S.K.P. critically reviewed and edited the manuscript. All authors read and approved the final version of the manuscript. All the authors agree to publish the manuscript.

## Acknowledgements

The authors gratefully acknowledge Yenepoya (Deemed to be University) for providing the infrastructure and state-of-the-art mass spectrometry facility required to conduct this study exclusively within our institution. We also acknowledge the support provided by the Department of Biotechnology (DBT) through the National Facility grant under the project “Skill Development in Mass Spectrometry-based Metabolomics Technology BIC” (BT/PR40202/BTIS/137/53/2023).

## Abbreviations

ACN: Acetonitrile
BCA: Bicinchoninic acid
CID: Collision-induced dissociation
CDK: Cyclin-dependent kinase
DDA: Data-dependent acquisition
DTT: Dithiothreitol
EDTA: Ethylenediaminetetraacetic acid
ERK: Extracellular signal-regulated kinase
FA: Formic acid
Fe-NTA: Ferric nitrilotriacetic acid
FR: Enriched and fractionated sample
FT: Final flowthrough
GO: Gene Ontology
HAT: Histone acetyltransferase
IAA: Iodoacetamide
IAP: Immunoaffinity purification
IMAC: Immobilized metal affinity chromatography
LC–MS/MS: Liquid chromatography tandem mass spectrometry
LFQ: Label-free quantification
MAPK: Mitogen-activated protein kinase
MS: Mass spectrometry
MS/MS: Tandem mass spectrometry
PBS: Phosphate-buffered saline
PTM: Post-translational modification
pY: Phosphotyrosine
RT: Room temperature
SDS-PAGE: Sodium dodecyl sulfate polyacrylamide gel electrophoresis
SOP: Standard operating procedure
TEABC: Triethylammonium bicarbonate
TFA: Trifluoroacetic acid
TiO₂: Titanium dioxide
TMT: Tandem mass tag
UE: Unenriched sample
UF: Enriched but unfractionated sample

